# Selective Activation of Na_V_1.3 Restores Lymphatic Contractility in Age and Injury

**DOI:** 10.64898/2025.12.15.694435

**Authors:** Katarina J. Ruscic, Reetu Singh, Lingshan Liu, Marla Marquez, Johanna J. Rajotte, Pinji Lei, Olivia D. Miller, Brian J. Kinsman, Meghan J. O’Melia, Dhruv Singhal, James W. Baish, Leigh D. Plant, Timothy P. Padera

**Affiliations:** Department of Anesthesiology, Massachusetts General Hospital and Harvard Medical School, Mass General Brigham, Boston, MA, 02114, USA; Edwin L. Steele Laboratories, Department of Radiation Oncology, Massachusetts General Hospital Cancer Center, Massachusetts General Hospital and Harvard Medical School, Mass General Brigham, Boston, MA, 02114, USA; Department of Immunology, Roswell Park Comprehensive Cancer Center, Buffalo, NY, 14263, USA; Brown University, Providence, RI, 02912, USA; Beth Israel Deaconess Medical Center, Division of Plastic and Reconstructive Surgery, Department of Surgery, Boston, MA, 02215, USA; Bucknell University, Department of Biomedical Engineering, Lewisburg, PA, 17837, USA; Department of Pharmaceutical Sciences, School of Pharmacy and Pharmaceutical Sciences, Bouvé College of Health Sciences, Northeastern University, Boston, MA, 02115, USA; Center for Drug Discovery, Northeastern University, Boston, MA, 02115, USA

**Author notes:** Corresponding Author: Timothy P. Padera, Ph.D. Massachusetts General Hospital Radiation Oncology-Cox 737, 100 Blossom St., Boston, MA 02114. Author Contribution Acknowledgements: K.J.R., L.D.P., and T.P.P. conceived and designed the study. K.J.R., R.S., L.L., M.M., J.J.R., B.J.K. and M.J.O. performed experiments. K.J.R., R.S., L.L., M.M., J.J.R., and O.D.M. analyzed the results of experiments in the study. P.L. analyzed single cell sequencing data. D.S. collected human lymphatic tissue samples. K.J.R. wrote the original first manuscript draft. All co-authors edited and approved the manuscript.

## Abstract

**Background:** Intrinsic lymphatic contractility is essential for tissue fluid balance, immunity and organ function, yet no FDA-approved pharmacologic treatments specifically restore lymphatic contractility. Lymph is returned to the circulation by ion channel-driven cyclic contractions of collecting lymphatic vessels. Although voltage-gated sodium (Na_V_) channels drive cardiomyocyte excitability, their role in lymphatic muscle cell (LMC) physiology is not well defined. We identified Na_V_1.3, a Na_V_ channel historically viewed as developmentally restricted and limited in adult tissues, as unexpectedly and selectively expressed in adult lymphatic muscle but absent from heart, vascular smooth muscle, and mature brain. We tested whether selective Na_V_1.3 activation restores impaired lymphatic pumping in aging and radiation injury.

**Methods:** Na_V_1.3 expression in LMCs was confirmed through single-cell RNA sequencing analysis and immunostaining of mouse and human lymphatic vessels. Lymphatic contractility was quantified by *in vivo* fluorescence lymphangiography and interstitial fluid clearance was measured with a new bioluminescence assay. Na_V_1.3 function was assessed in young, aged, and radiation-injured mice. Na_V_1.3 knockout (*Scn3a^-/-^*) mice established the requirement of Na_V_1.3 for basal lymphatic excitability and responsiveness to the Na_V_1.3-specific activator, Tf2.

**Results:** In mouse and human lymphatic vessels, Na_V_1.3 is expressed in adult LMCs. Although dispensable for basal lymphatic contractions, Na_V_1.3 acted as a pharmacologically recruitable reserve that amplified contractile output. Acute Na_V_1.3 activation with Tf2 increased lymphangion ejection fraction and accelerated interstitial fluid clearance. Tf2 fully restored lymphatic pumping in aged mice and partially rescued radiation-induced contractile deficits. All Tf2 responses were abolished in *Scn3a^-/-^* mice, confirming Na_V_1.3 dependence.

**Conclusions:** Na_V_1.3 is a selectively druggable ion channel in adult lymphatic muscle that can be recruited to restore lymphatic pump function across aging and injury. Targeted Na_V_1.3 activation provides a molecular entry point for treating diseases characterized by lymphatic pump failure, a domain with no existing pharmacologic therapies.

## INTRODUCTION

The lymphatic system is a network of vessels and lymph nodes that maintains tissue fluid balance, supports immune cell trafficking, and aids lipid absorption [1–5]. Lymphatic function declines with age [6–11], after radiation injury [12–17], and contributes to pathological fluid imbalance in conditions such as cancer therapy-associated lymphedema [18–20], neurodegenerative diseases [10, 21–24], inflammatory diseases [25–27], and cardiovascular diseases such as heart failure, hypertension and atherosclerosis [28–31]. Collecting lymphatic vessels are invested in lymphatic muscle cells (LMCs) [7] and actively propel fluid through phasic contractions of lymphangions, which are vessel segments bound by unidirectional valve [32–35]. Despite the critical role of intrinsic lymphatic pumping in fluid homeostasis, treatments for lymphatic dysfunction do not target lymphatic contractility directly and are limited to mechanical interventions such as compression, massage, or surgical rerouting of lymph flow [18, 36–39]. There are currently no FDA-approved medications indicated to specifically target lymphatic contractile function [40].

Contraction of LMCs relies on the generation of an ion channel mediated action potential [34, 35]. Excitable tissues such as the heart, brain, and skeletal muscle rely on voltage-gated ion channels to initiate and coordinate electrical signaling. In particular, voltage-gated sodium (Na_V_) channels pass the initial depolarizing current in these excitable tissues and are extensively targeted in clinical practice to modulate cardiac arrhythmias, epilepsy, and pain [41–43]. In lymphatic contraction, tetrodotoxin-sensitive sodium currents have been reported in sheep lymphatic vessels [44] and Na_V_1.3 has been found in human lymphatic vessels [45]. However, further study on the role of Na_V_ channels in regulating lymphatic contractility is needed to explore their therapeutic potential.

Our analysis of single-cell RNA sequencing data from murine collecting lymphatics in multiple studies revealed that *Scn3a*, the gene encoding Na_V_1.3, is selectively present in LMCs [7, 46]. Unlike other Na_V_ isoforms, Na_V_1.3 is absent from cardiac and neuronal tissues after development [47], making it a promising candidate for selective pharmacologic modulation of lymphatic excitability in adults. Historically, Na_V_1.3 has been viewed as a transient, developmentally-restricted channel, highly expressed in the embryonic nervous system but largely silenced in adult physiology [47–50], other than expression after neuronal injury [43], in some neutrophils [51], and in adrenal chromaffin cells [52]. Its presence as a dominant isoform in adult LMCs introduces an unexpected and exciting new role for Na_V_1.3 in adult physiology. While multiple membrane proteins are important in lymphatic contractility, including L-type calcium channels [53–55], TMEM16A (Ano1) [56], and several voltage-gated potassium channels [57], these ion channels are widely expressed in many tissues, such as the heart, brain, and blood vasculature, raising concerns about systemic pharmacological effects when targeted [58]. The restricted expression of Na_V_1.3 in adult mice to LMCs [7] offers a unique and potentially safe avenue for modulating lymphatic function in a targeted way. Prior work has demonstrated Na_V_-dependent electrical activity in lymphatic vessels across species [44, 45], with Na_V_1.3 as the primary human isoform [45]. However, the therapeutic potential of specifically activating Na_V_1.3 to restore or enhance lymphatic function has not been investigated.

We hypothesized that Na_V_1.3 is a key regulator of lymphatic contractility and that its activation could enhance lymphatic pumping and fluid clearance. To test this, we performed *in vivo* fluorescence lymphangiography and bioluminescence clearance assays in mice exposed to Tf2, a specific Na_V_1.3 activator peptide originally isolated from *Tityus fasciolatus* scorpion venom [59]. Tf2 significantly increased lymphatic ejection fraction and this translated into accelerated interstitial fluid clearance, confirming the functional relevance of Na_V_1.3 activation. Finally, we evaluated the therapeutic potential of Na_V_1.3 activation in models of age- and radiation-induced lymphatic dysfunction [6, 7, 12, 18], where Tf2 helped restore contractile function. These findings highlight Na_V_1.3 as a lymphatic-specific therapeutic target with the potential to restore lymphatic function in clinical conditions characterized by failure of lymphatic contractility, which currently have no targeted pharmacologic treatments available.

## METHODS

### Data Availability

All detailed experimental protocols are provided in the Supplemental Material. Materials and reagents will be made available upon reasonable request.

### Human Tissue

Human axillary lymphatic vessels were collected intraoperatively from patients undergoing lymphatic reconstruction at Beth Israel Deaconess Medical Center under an Institutional Review Board approved protocol, with written informed consent. Free vessel ends that would otherwise be discarded were placed on ice immediately, embedded in OCT (Tissue-Tek), and cooled in an ethanol slurry within 10 minutes of excision, then stored at −80 °C until use for immunofluorescence.

### Mice and Models

All animal studies were performed under protocols approved by the Massachusetts General Hospital Institutional Animal Care and Use Committee. Wild-type C57BL/6 mice were studied at young adult (8–12 weeks), 12 months, and 18 months of age to model physiologic aging. *Scn3a*⁻*/*⁻ mice and Prox1-GFP reporter mice were used for functional and imaging studies. Radiation-induced lymphatic injury was established by localized hindlimb irradiation using fractionated X-ray exposure to a cumulative dose of 18 Gy, delivered over three days to minimize acute toxicity. Irradiation was performed using a Precision X-ray irradiator (225 kVp, 13.30 mA; dose rate 1.98 Gy/min).

### Molecular Analyses

*Scn3a* expression in LMCs was assessed by integrating published single-cell RNA-sequencing datasets (GSE248658 and GSE277843), quantitative PCR of pooled inguinal axillary collecting lymphatic vessels, and immunofluorescence staining of murine and human lymphatic tissue. Human samples were analyzed as frozen tissue sections, and mouse lymphatic vessels were prepared as whole mounted lymphatic vessels. Tissues were stained with antibodies against α-SMA and Na_V_1.3, with DAPI nuclear counterstaining. Confocal fluorescence imaging was performed on a Zeiss LSM 880 to confirm cellular localization of Na_V_1.3 in LMCs. Additional details on antibody sources and staining protocols are provided in the Supplemental Material.

### *In Vivo* Lymphatic Function

Collecting lymphatic contractility was quantified *in vivo* by intravital epifluorescence lymphangiography. After induction of anesthesia, mice received an intradermal injection of 3 μL of 2% FITC–dextran (2 MDa; Millipore Sigma) into the dorsal hind footpad and afferent collecting vessels were surgically exposed. The hindlimb was immersed in a static pool of 150 μL sterile 0.9% saline (vehicle), which was replaced with test solution when indicated. Time-lapse epifluorescence imaging was performed on a microscope (Olympus IX70) equipped with a camera (Hamamatsu) for 60–120 seconds at ∼7.5 frames per second. Vessel diameters were extracted frame-by-frame with custom MATLAB scripts to calculate mean, systolic, and diastolic diameters, contraction frequency, and ejection fraction.

Interstitial fluid clearance was assessed by bioluminescence. After anesthesia and surgical exposure of the hindlimb collecting vessel draining to the popliteal node, mice received intraperitoneal D-luciferin (500 μL of 30 mg/mL; PerkinElmer). The exposed hindlimbs were then immersed in sterile 0.9% saline or 10 μM Tf2. Luciferase (2 μL of 5 mg/mL; PerkinElmer) was injected intradermally into each footpad, and serial imaging on the IVIS Spectrum System quantified signal decay at the injection site. Clearance kinetics were analyzed with a two-compartment model in MATLAB.

### Statistics

Data are expressed as mean ± SEM. Repeat measures one-way ANOVA with Dunnett’s multiple-comparison test was used for sequential measurements within the same animals with an alpha of 0.05, and adjusted P values with * significance for P<0.03, ** P<0.002, *** P<0.0002. Unpaired t-tests were used for comparisons between independent groups with the same significance cutoffs as above. Comparisons within the same animal were performed using a two-tailed paired t-test, with * indicating P<0.05. Analyses were conducted in GraphPad Prism.

## RESULTS

### Na_V_1.3 is present on human and mouse lymphatic muscle cells

As Na_V_1.3 has been previously shown on sheep and human collecting lymphatic vessels [45], we analyzed published murine single-cell RNA sequencing datasets [7] [46] (Fig. 1A). The data showed expression of *Scn3a*, which encodes the voltage-gated sodium channel Na_V_1.3, on *Acta2+Myhll+* LMCs. Immunofluorescence staining of a human axillary lymphatic vessel obtained from a lymphatic reconstruction surgery confirmed the presence of Na_V_1.3 in α-smooth muscle actin (α-SMA) positive LMCs (Fig. 1B). Although both markers are present in the same cells, their subcellular distributions are distinct with Na_V_1.3 being a membrane-bound protein and α-SMA present in the cytoplasm (Fig. 1B inset shows higher magnification of the boxed region to highlight localization of Na_V_1.3 to LMCs). Similar expression patterns were observed in whole mount preparations of mouse inguinal axillary lymphatic vessels, where Na_V_1.3 was detected in α-SMA-positive LMCs along the vessel wall (Fig. 1C). Prox1 is a transcription factor that defines lymphatic endothelial cell (LEC) identity and is used as a marker to distinguish LECs from surrounding non-endothelial cells [60, 61]. Staining in Prox1-GFP mice [61] demonstrated Na_V_1.3 localization to LMCs, distinct from the GFP-positive Prox-1 positive LECs (Fig. 1C). Our data show that only a portion of the murine LMCs stain positive for Na_V_1.3, consistent with our single cell sequencing dataset ([7]), in which approximately one-third of LMCs expressed *Scn3a* mRNA, suggesting heterogeneity in Na_V_1.3 expression in mouse LMCs.

**Figure 1:**
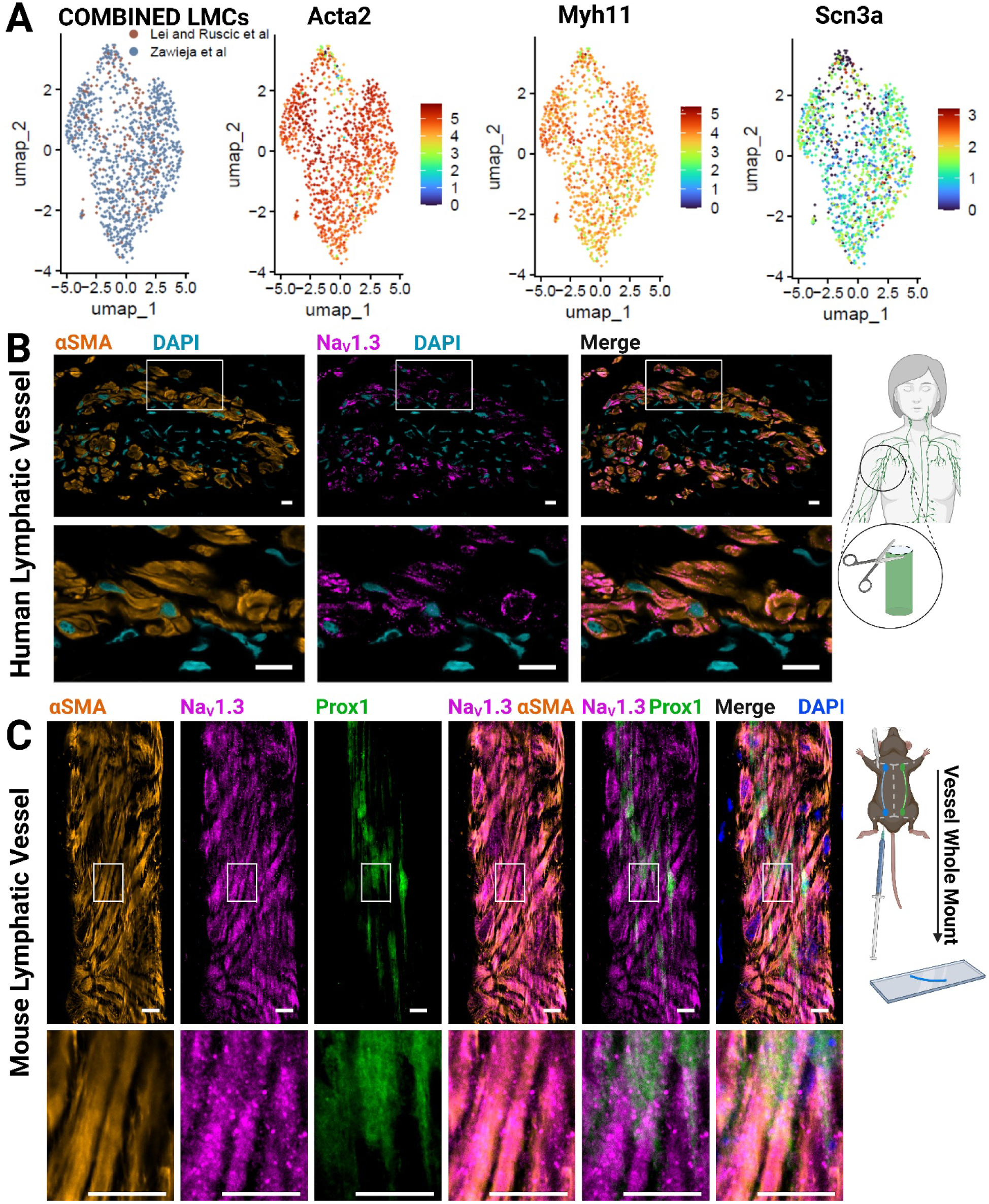
Na_V_1.3 protein is present in lymphatic muscle cells of humans and mice. (A) UMAP plot of 1,196 LMCs from published single-cell RNA-seq datasets GSE248658 [7] and GSE277843 [46]. Left to right, colored by: dataset, *Acta2* expression, *Myh11* expression, and *Scn3a* expression. Gene expression levels are log-normalized and scaled. (B) Immunofluorescence for α-SMA (orange), Na_V_1.3 (magenta), and DAPI (cyan) of a human axillary lymphatic vessel collected intraoperatively. Scale bar is 10 µm. (C) Whole mount of mouse inguinal axillary lymphatic vessel stained for α-SMA (orange), Na_V_1.3 (magenta), and DAPI (cyan) in a Prox1-GFP transgenic mouse (Prox1, green) (Scale bar, 10 µm). For panels B and C, the second row shows magnified insets of the boxed regions in the first row.

### Activation of Na_V_1.3 enhances lymphatic contractility, but Na_V_1.3 is not necessary for phasic lymphatic contractions

To determine if Na_V_1.3 plays a functional role in lymphatic contraction, we assessed collecting lymphatic vessel function *in vivo* using epifluorescence lymphangiography as previously described [62] and measured lymphatic vessel diameter changes over time in collecting lymphatic vessels draining to the popliteal lymph node (Fig. 2A, Supplemental Video 1). In order to evaluate the potential of Na_V_ channels to modulate lymphatic contraction, we tested the pan-Na_V_ agonist, veratridine, which prolongs the open state of all Na_V_ isoforms. Veratridine exposure results in sustained cell depolarization by preventing fast inactivation of the sodium channel [63] and has previously been shown to induce contractions and increase tension in human thoracic duct and mesenteric lymphatic ring segments in a concentration-dependent manner [45]. In young adult wild-type mice (8–12 weeks old), veratridine dose-dependently enhanced lymphatic contractility (Fig. 2B, Supplemental Video 2) as quantified by ejection fraction, which reflects the strength of lymphangion contraction [34, 62, 64]. Mean, systolic, and diastolic diameters were measured within each lymphangion (Fig. 2A). The increase in ejection fraction is related to a decrease in systolic diameter with increasing veratridine concentration with a mean systolic diameter of 33 µm in 10 µM veratridine compared to 40 µm in the saline control (p < 0.05; Fig. 2B, Supplemental Video 2).

**Figure 2:**
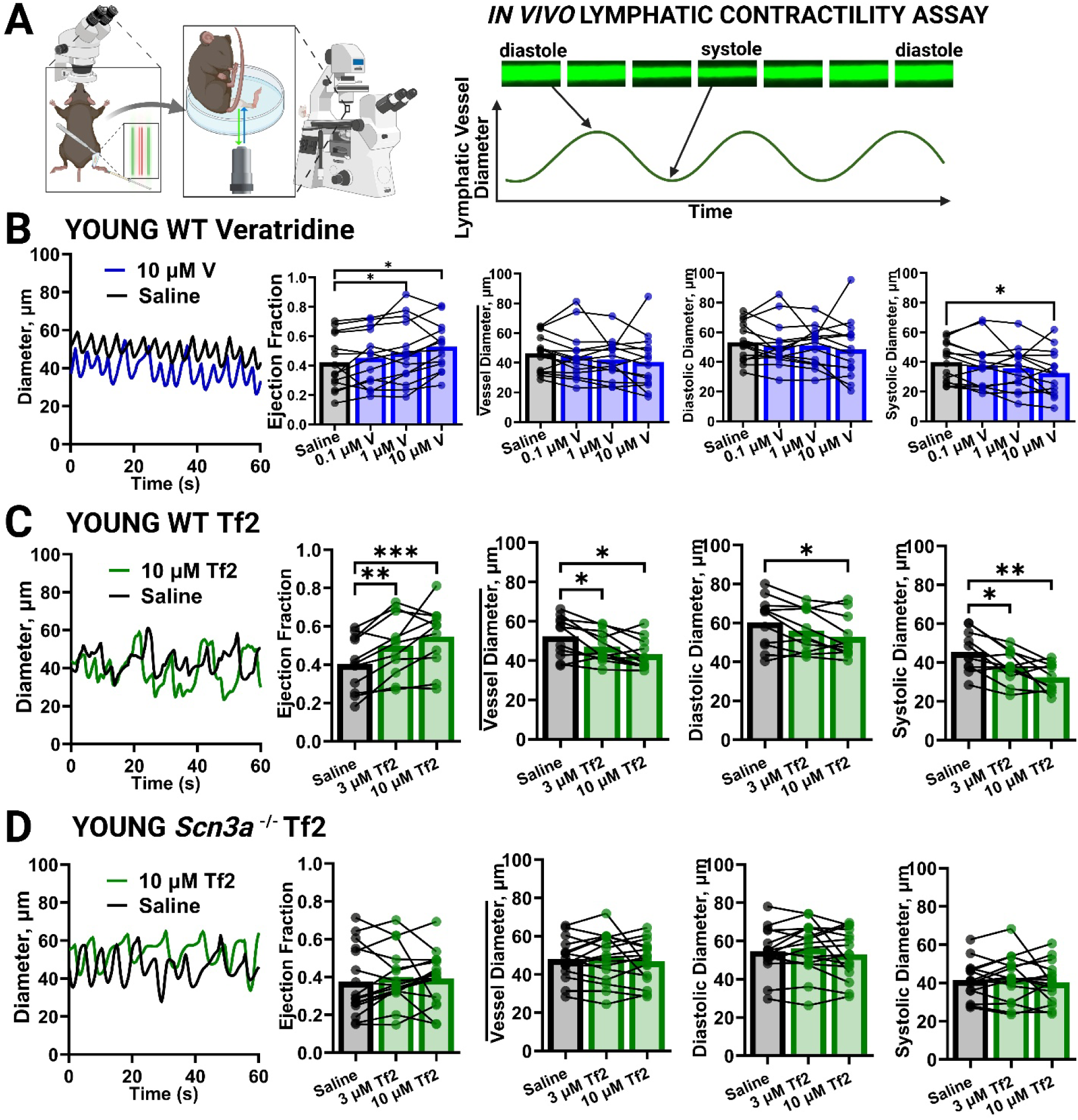
Activation of Na_V_1.3 enhances lymphatic contractility, but Na_V_1.3 is not necessary for lymphatic pumping. (A) *In vivo* lymphatic contractility of the collecting lymphatic vessels to the popliteal lymph node was measured using epifluorescence lymphangiography. (B) In young (8-12-week-old) wild-type mice, a representative trace of diameter (µm) vs. time (seconds) is shown with the hindlimb exposed to saline (vehicle, black) or 10 µM veratridine (V, blue). The ejection fraction shows the strength of contraction in the same lymphangion exposed to saline (vehicle, black) and increasing concentrations of veratridine (V, blue): 0.1 µM, 1 µM, and 10 µM. The mean vessel diameter (*vessel diameter*, µm), systolic diameter (µm), and diastolic diameter (µm) are shown for the same four drug conditions. N = 14 mice. (C,D) In young wild-type mice (C) and young *Scn3a-/-* mice (D), a representative trace of diameter (µm) vs. time (seconds) is shown with the hindlimb exposed to saline (vehicle, black), 3 µM Tf2, or 10 µM Tf2 (green). Ejection fraction, *vessel diameter* (µm), systolic diameter (µm), and diastolic diameter (µm) are shown in the same lymphangion sequentially exposed to saline (vehicle, black), 3 µM Tf2, or 10 µM Tf2 (green). N = 11 wild-type young mice, 16 *Scn3a-/-* young mice. Repeat measures one-way ANOVA with Dunnett’s multiple-comparison test was used for sequential measurements within the same animals with an alpha of 0.05, and adjusted P values with * significance for P<0.03, ** P<0.002, *** P<0.0002.

To more specifically target Na_V_1.3, we used the Na_V_1.3-selective activator, Tf2. Originally identified from the venom of the *Tityus fasciolatus* scorpion from Brazil, Tf2 is a 64-amino acid β-NaScTx peptide that left-shifts the activation voltage threshold of Na_V_1.3, thus increasing the probability of the channel being open at the resting membrane potential [59, 65]. In young adult wild-type mice, lymphatic contractility was increased by Tf2 exposure (Fig. 2C, Supplemental Video 3). Representative diameter tracings over time demonstrate rhythmic contractions under baseline conditions (saline) with a dose-dependent increase in the ejection fraction following sequential exposure to saline, 3 µM Tf2, and 10 µM Tf2 (Fig. 2C). This was accompanied by a concomitant decrease in the overall mean vessel diameter with increasing concentrations of Tf2, driven by a strong decrease in the systolic diameter and a mild decrease in the diastolic diameter (Fig. 2C).

*Scn3a*^⁻^*^/^*^⁻^ mice, which lack Na_V_1.3, showed no change in contractile parameters following exposure to any concentration of Tf2 (Fig. 2D, Supplemental Video 4), indicating that the lymphatic response to Tf2 is dependent on Na_V_1.3. Surprisingly, *Scn3a*^⁻^*^/^*^⁻^ mice exhibited regular, rhythmic phasic contractions under baseline conditions, with no evidence of lymphatic dysfunction or overt edema. Baseline contractile parameters, including ejection fraction, mean diameter, and systolic/diastolic diameters, were comparable to those observed in wild-type mice (Fig. 2D).

We observed a small decrease in contraction frequency over time during intravital imaging, even when only saline was applied (Supplemental Fig. 1). Accordingly, we interpret changes in frequency with caution and present them in Supplemental Fig. 1.

### TTX does not abolish lymphatic contractility but prevents Tf2-enhanced lymphatic contraction

To further assess whether the contractile-enhancing effects of Tf2 on lymphatic vessels are dependent on Na_V_ channel activity, we examined the impact of tetrodotoxin (TTX)—a potent blocker of a subset of Na_V_ channels that includes Na_V_1.1, Na_V_1.2, Na_V_1.3, Na_V_1.4, Na_V_1.6, and Na_V_1.7—on lymphatic vessel contractility *in vivo* (Fig. 3A). Using epifluorescence lymphangiography, we first measured the contractile behavior of collecting lymphatic vessels draining to the popliteal lymph node in young wild-type mice under sequential exposure to saline (vehicle), 10 µM Tf2, and 1 µM TTX. Tf2 increased the lymphangion ejection fraction compared to the saline baseline in most animals, consistent with previous experiments (Fig. 3B). TTX exposure then abolished the gain in ejection fraction caused by Tf2 (Fig. 3B). The systolic diameter likewise decreased with Tf2 exposure, and then increased back to baseline with TTX exposure (Fig. 3B). We next changed the order of exposure to TTX and Tf2, first exposing the vessel to saline (vehicle) then 1 µM TTX followed by 10 µM Tf2 (Fig. 3C). The exposure of lymphatic vessels to TTX from a saline baseline did not change the baseline ejection fraction or any lymphatic vessel diameter measurements in most animals, suggesting that the lymphatic contractility was not dependent on a TTX-sensitive Na_V_ channel. These data are consistent with the results from *Scn3A^-/-^* mice (Fig. 2D). The mice in Fig. 3C were then exposed to Tf2 after exposure to TTX without washout of the TTX. Under those conditions of pre-blockade with TTX, Tf2 did not significantly increase the ejection fraction (Fig. 3C). Tf2 likewise did not significantly change mean vessel diameter, systolic diameter, or diastolic diameter when applied after exposure to TTX (Fig. 3C). Hence, our data show that when the pores of Na_V_1.3 channels are tightly blocked by TTX, the Tf2-induced left-shift in voltage-dependent activation does not enhance contractility (Fig. 3C).

**Figure 3:**
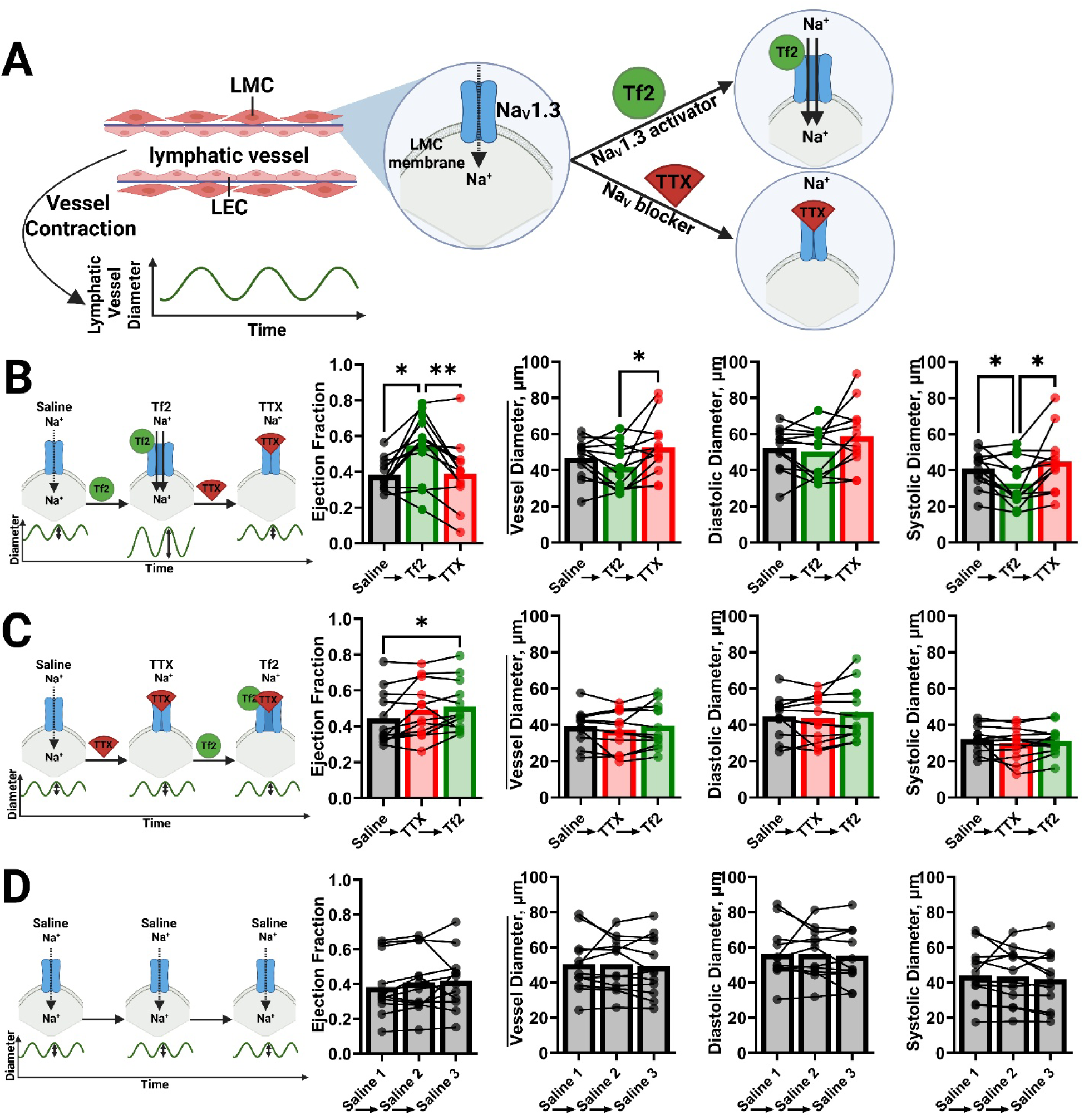
TTX does not abolish lymphatic contractility but prevents Tf2-enhanced lymphatic contraction. (A) *In vivo* lymphatic contractility of the collecting lymphatic vessel to the popliteal lymph node was measured with epifluorescence lymphangiography in young wild-type mice with the vessel bathed in saline (vehicle, black), 10 µM Tf2 (activator, green), or 1 µM TTX (blocker, red). In (B), lymphatic contractility [ejection fraction, *vessel diameter* (µm), systolic diameter (µm), and diastolic diameter (µm] was first measured with the hindlimb in a saline (vehicle, black) to establish baseline activity. The same lymphangion was sequentially incubated with 10 µM Tf2 (green) followed by incubation with 1 µM TTX (red), with a solution change performed between each treatment. In (C), lymphatic contractility is measured in saline (vehicle, black), followed by 1 µM TTX (red), and then exposed to 10 µM Tf2 (green). In (D), lymphatic contractility is measured in saline vehicle (saline 1, black), followed by a solution change with saline (saline 2, black), and a third solution change with saline (saline 3, black). N=12 wild-type young mice per group. Repeat measures one-way ANOVA with Dunnett’s multiple-comparison test was used for sequential measurements within the same animals with an alpha of 0.05, and adjusted P values with * significance for P<0.03, ** P<0.002, *** P<0.0002.

Because *in vivo* lymphatic physiological experiments may be affected by perturbations such as solution changes, and because the behavior of the lymphatic vessel may change with time (Supplemental Fig. 1), we performed saline-only solution changes that mirrored the timing of experimeints in Figs. 3B and 3C. The resultant saline control data show that the ejection fraction, mean vessel diameter, diastolic diameter and systolic diameter measurements do not change between saline changes (Fig. 3D). Thus the effects in Figs. 3B and 3C are not artifacts of solution changes and incubation time.

### Activation of Na_V_1.3 by Tf2 speeds the clearance of interstitial fluid

To determine whether activation of Na_V_1.3 channels enhances lymphatic-mediated clearance of interstitial fluid, we developed a bioluminescence-based assay in young mice. After removing the skin and soft tissue from the posterior hindlimbs to expose the collecting lymphatic vessels to the popliteal lymph node, mice received an intraperitoneal injection of D-luciferin to systemically distribute this substrate into all tissues after incubation. A subcutaneous injection of 2 μL luciferase into the dorsum of each footpad delivered the enzyme only into a localized region, serving as a controlled interstitial fluid bolus (Fig. 4A, left). The localized luciferin/luciferase reaction in the footpads allowed us to track the function of the lymphatic vessels draining the footpad and clearing away the luciferase interstitial fluid bolus. The underside of each hindlimb in the supine mouse was then exposed to saline (vehicle control) or 10 µM Tf2 (Fig. 4A). Each footpad emits photons (Fig. 4A, right), which then decreases as the interstitial fluid bolus of luciferase is cleared from the footpads by the exposed lymphatic vessels (Fig. 4B).

**Figure 4:**
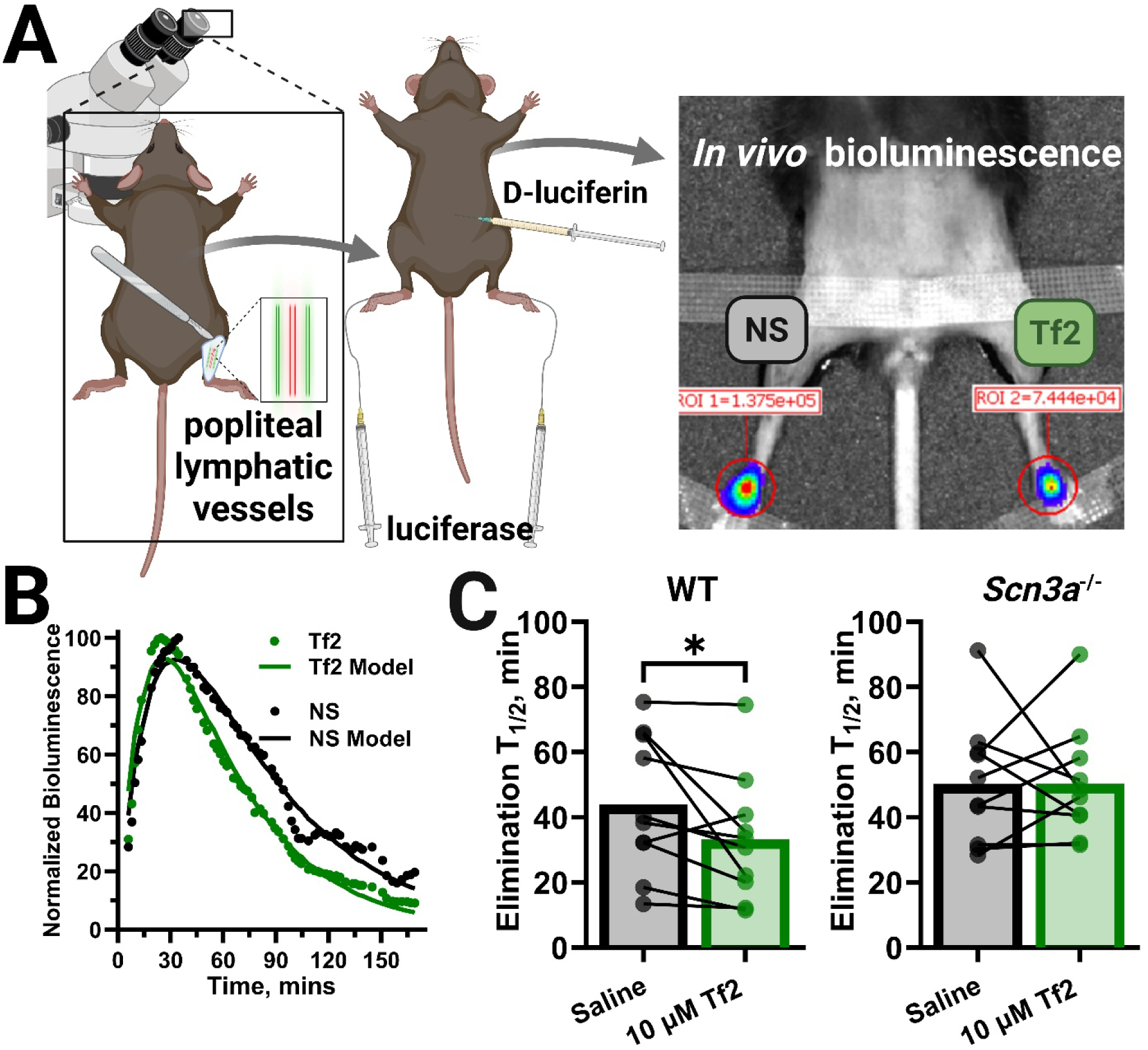
Activation of Na_V_1.3 by Tf2 speeds clearance of interstitial fluid. (A) Young (8-12 week) mice undergo surgical removal of the skin and soft tissue covering the posterior hindlimb bilaterally, followed by intraperitoneal injection of 500 µL of 30 mg/mL D-luciferin. 2 µL of 5 mg/mL luciferase is then injected into the dorsum of both footpads. Bioluminescence signal is collected over time in bilateral footpads with the underside of the hindlimb exposed to either 150 µL of saline or 150 µL of 10 µM Tf2. (B) Representative example of normalized bioluminescence signal as a function of time (minutes) in a mouse with one hindlimb exposed to saline (vehicle, black dots) and the other hindlimb exposed to Tf2 (green dots). Extracellular fluid elimination time constants are modeled from a two-compartment model (black and green curves). (C) Fluid elimination half-life (T_1/2_) for young wild-type mice (N = 10 mice) and young *Scn3a- /-* mice (N = 10 mice) in paired saline (vehicle, black) and Tf2-exposed hindlimbs (green). Comparisons within the same animal were performed using a two-tailed paired t-test, with * indicating P<0.05.

Representative time-course data from a wild-type mouse reveals faster decay of the bioluminescence signal in the Tf2-exposed limb compared to the saline-exposed control (Fig. 4B). Kinetic modeling using a two-compartment model showed an increased fluid clearance rate constant in the Tf2 condition, indicating more rapid clearance of luciferase in the Tf2-exposed hindlimb compared to the saline-exposed hindlimb.

Across animals (N=10 per group), Tf2 significantly increased the luciferase clearance rate in young wild-type mice (Fig. 4C, left). In contrast, young *Scn3a*^⁻^*^/^*^⁻^ mice showed no overall difference in luciferase clearance between Tf2- and saline-exposed limbs (Fig. 4C, right), demonstrating that the effect of Tf2 on luciferase clearance is dependent on Na_V_1.3.

### Activation of Na_V_1.3 by Tf2 in aged mice rescues the age-related decrease in lymphatic contractility

Lymphatic contractility declines with age [6, 7, 66], as demonstrated by reduced ejection fraction in older animals compared to young animals (mean ejection fraction in saline of 0.40±0.04 in young vs. 0.27±0.03 in 12 month old and 0.28±0.04 in 18 month old mice, p<0.05). To determine whether this age-related decline is associated with changes in Na_V_1.3 expression, we analyzed *Scn3a* transcript levels in LMCs across the murine lifespan. Single-cell RNA sequencing combining our results and those of Zawieja et al. [7, 46] revealed preservation of *Scn3a* gene expression from 6-week-old mice, adult mice, and 12- and 18-month-old mice (Fig. 5A). mRNA expression, measured by quantitative PCR of whole lymphatic vessels, demonstrated a relative reduction in *Scn3a* expression in aged mouse vs. young mouse whole vessels, consistent with a known reduction in lymphatic vessel coverage by LMCs with age [7, 67]. These data suggest that *Scn3a* remains expressed on a per LMC basis in older animals. Embryonic brain served as a positive control tissue for *Scn3a* expression to which test samples were normalized [47–50]. *Scn3a* expression was negligible in young *Scn3a*^⁻^*^/^*^⁻^ mice (Fig. 5B), confirming these animals as full knockouts [52, 68].

**Figure 5:**
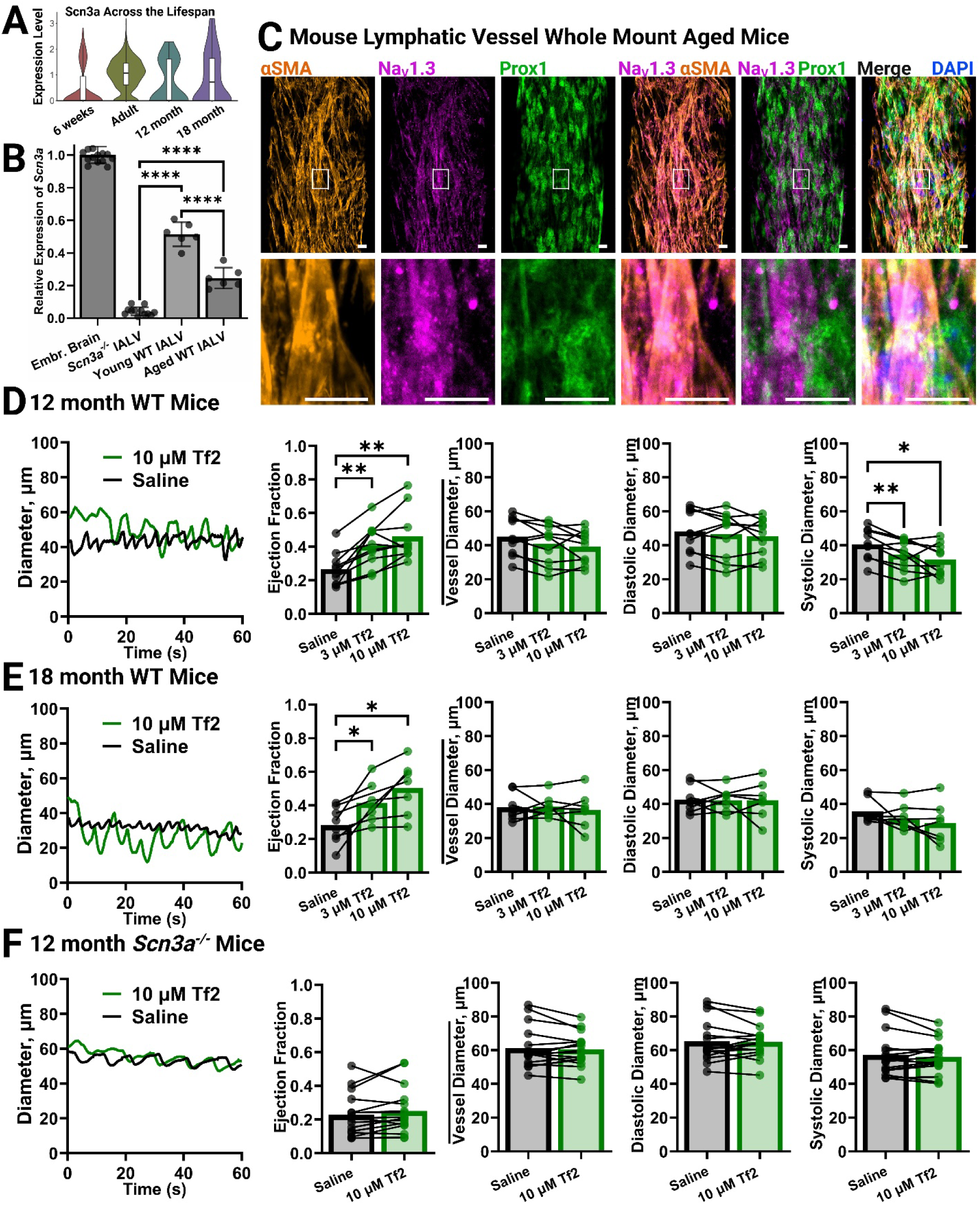
Activation of Na_V_1.3 by Tf2 in aged mice rescues lymphatic contractility. (A) Violin plots showing the single-cell expression level of *Scn3a* in LMCs, grouped by age, from published datasets GSE248658 [7] and GSE277843 [46]. Gene expression levels are log-normalized and scaled. (B) Quantitative PCR expression of *Scn3a* in whole inguinal axillary lymphatic vessels from young (8-12 week) wild-type mice (N=8 vessels, 4 mice), aged (18-23 month) wild-type mice (N=8 vessels, 4 mice), and young (8-12 week) *Scn3a*-/- mice (N=16 vessels, 8 mice) relative to expression of *Scn3a* in embryonic brain (2-4 pooled PCR triplicate runs). (C) Whole mount of 18-month old mouse inguinal axillary lymphatic vessel stained for α-SMA (orange), Na_V_1.3 (magenta), and DAPI (cyan) in a Prox1-GFP mouse (Prox1, green) (Scale bar, 10 µm, second row shows magnified insets of boxed regions in the first row). (D-F) In vivo lymphatic contractility of the collecting lymphatic vessel to the popliteal lymph node was measured using epifluorescence lymphangiography in (D) 12 months old wild-type, (E) 18 month old wild-type and (F) 12 month old *Scn3a-/-* mice. A representative trace of diameter (µm) vs. time (seconds) is shown with the hindlimb exposed to saline (vehicle, black) or 10 µM Tf2 (green). The ejection fraction shows the strength of contraction in the same lymphangions exposed to saline (vehicle, black) followed by 3 µM, and 10 µM Tf2 (green). The mean vessel diameter (*vessel diameter*, µm), systolic diameter (µm), and diastolic diameter (µm) are shown for the same three drug conditions. N = 10 mice for 12 month old wild-type. N = 8 mice for 18 month old wild-type, N = 16 mice for 12 month old *Scn3a-/-* mice. Repeat measures one-way ANOVA with Dunnett’s multiple-comparison test was used for sequential measurements within the same animals with an alpha of 0.05, and adjusted P values with * significance for P<0.03, ** P<0.002, *** P<0.0002.

Analysis of inguinal axillary lymphatic vessels from 18 month old Prox1-GFP mice (expressing GFP under the lymphatic endothelial cell transcription factor *Prox1*, green [61]) confirmed the presence of Na_V_1.3 protein (magenta) in LMCs marked by α-SMA (orange), with nuclear counterstaining by DAPI (blue). Na_V_1.3 localizes to LMCs within the vessel (Fig. 5C), showing it is potentially a viable pharmacological target in 18 month old mice.

To test whether pharmacologic activation of Na_V_1.3 could restore contractility in aged lymphatic vessels, we exposed middle-aged (12 month old) and very aged (18 month old) mice to Tf2. In vivo lymphangiography revealed that 10 µM Tf2 increased the ejection fraction in 12-month old mice by 70% (p<0.05), and in 18 month old mice by 90% (p<0.05) (Fig. 5D, Fig. 5E). In 12-month old mice, the systolic diameter significantly decreased in a dose-dependent manner with increasing concentrations of Tf2. In 18-month old mice, 3 and 10 µM Tf2 significantly increase EF compared to saline baseline in the same mice (p<0.05) (Fig. 5E). Notably, EF in 18 month old mice treated with 10 µM Tf2 (0.50) was not significantly different from EF in young mice at baseline (0.40, p=0.19), consistent with a reversal of age-related decline in lymphatic contractility by Na_V_1.3 activation (Fig. 2C). In contrast, aged *Scn3a*⁻*/*⁻ mice exhibited no response to 10 µM Tf2, confirming that the restorative effect of Tf2 on lymphatic contractility is dependent on the presence of functional Na_V_1.3 channels (Fig. 5F). These results suggest that Tf2 can rescue age-associated lymphatic dysfunction, in an Na_V_1.3 dependent manner.

### Activation of Na_V_1.3 partially rescues radiation-induced decreased lymphatic contractility

X-ray radiation is known to decrease lymphatic contractility [12], impair lymphatic endothelial barrier function [13], change lymph vessel caliber [14], and inhibit lymph drainage in edema models [15], amongst other unfavorable effects [16, 17]. We developed a model of radiation-induced hindlimb lymphatic injury with similar total radiation dose as prior murine radiation studies [17, 69] divided into three irradiation exposures to minimize the toxicity of each session. Mice were exposed to radiation or sham-treatment to the right hindlimb over three consecutive days (6 Gy per day) for a cumulative dose of 0 or 18 Gy X-rays (Fig. 6A). Mice receiving 18 Gy radiation showed decreased ejection fraction (Fig. 6B) and contraction frequency (Fig. 6C) in the collecting lymphatic vessel to the popliteal lymph node compared to the sham-radiation control group. Subsequent administration of 10 µM Tf2 to the hindlimb significantly increased the ejection fraction of the 18 Gy irradiated mice (Fig. 6B), indicating partial rescue of lymphatic vessel contractility. Tf2 did not rescue the decrease in contraction frequency (Fig. 6C). These findings suggest that activation of Na_V_1.3 by Tf2 can partially restore lymphatic contractile function after radiation-induced impairment.

**Figure 6:**
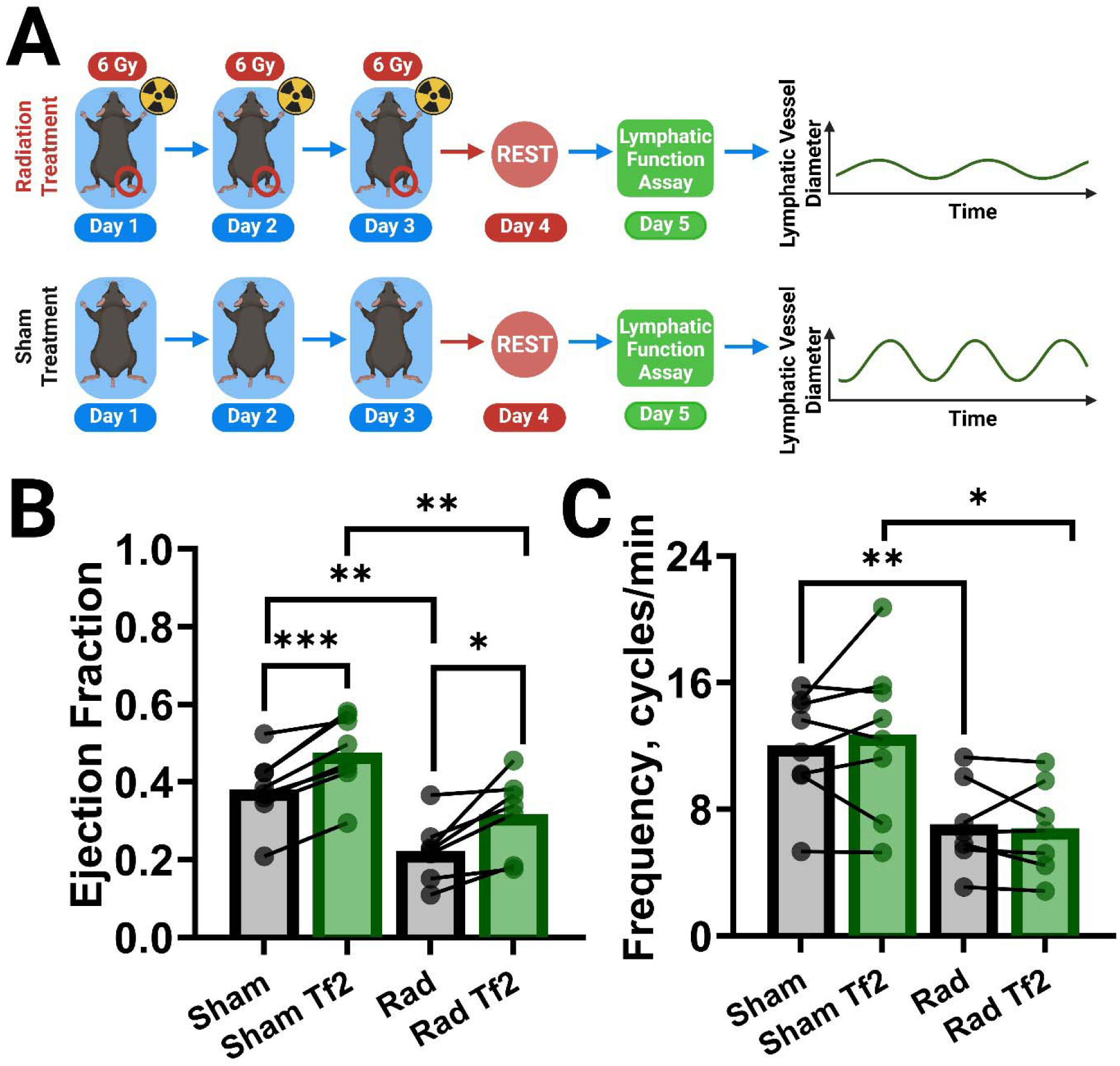
X-ray radiation decreases lymphatic contractility and Tf2 partially rescues contractility. (A) Young wild-type mice were assigned to either a sham or radiation treatment group and were exposed to 0 Gy or 6 Gy per day to the right hindlimb for three consecutive days. On day 5, mice underwent *in vivo* epifluorescence lymphangiography. (B) Ejection fraction and (C) contraction frequency of lymphangions in sham and radiation-treated groups under saline (vehicle, black) and Tf2 (green) for N = 7-8 mice per group. Repeat measures one-way ANOVA with Dunnett’s multiple-comparison test was used for sequential measurements within the same animals with an alpha of 0.05, and adjusted P values with * significance for P<0.03, ** P<0.002, *** P<0.0002. Unpaired t-tests were used for comparisons between independent groups with the same significance cutoffs

## DISCUSSION

Na_V_1.3 has long been regarded as a transient, developmentally restricted ion channel, prominent in the embryonic nervous system but largely absent in mature tissues [47–50]. Our data show that Na_V_1.3 is found on LMCs in adults, making it an attractive and potentially selective target to modify lymphatic contraction. In this study, we show that activation of Na_V_1.3 by the isoform-selective modulator Tf2 [59] enhanced lymphatic contractility and accelerated tissue clearance of a fiducial dye in young mice as well as partially rescued age-associated lymphatic dysfunction and radiation-induced lymphatic injury. These findings position Na_V_1.3 as a tissue-specific therapeutic target for disorders of impaired lymphatic pumping, which currently lack pharmacologic treatments.

We found that Na_V_1.3 is not required for baseline lymphatic excitability, as *Scn3a*⁻*/*⁻ mice retain rhythmic spontaneous contractions. However, targeted activation of Na_V_1.3 with Tf2 increases contraction strength and interstitial fluid drainage. These effects were completely abolished in *Scn3a*-deficient animals, confirming that Tf2 acts specifically through Na_V_1.3 to enhance lymphatic contractility. Further, Tf2-mediated increases in lymphatic contractility were precluded by the Na_V_ blocker TTX, confirming a Na_V_ target of Tf2. Unlike previously described ion channels in lymphatics [34, 35], including L-and T-type calcium channels [53–55, 70–72], TMEM16A (Ano1) [56], and voltage-gated potassium channels [57], Na_V_1.3 is not broadly expressed across excitable tissues [47–50], minimizing the risk of systemic off-target effects if pharmacologically activated.

Aging is a natural biological process causing decreased lymphatic function, and radiation can impair lymphatic contractility after cancer therapy and predispose patients to lymphedema [6–17]. Using aging and radiation treatment as models of lymphatic impairment, we observed depressed contractility that was partially rescued by pharmacologic activation of Na_V_1.3. Notably, in 18-month-old mice, Tf2 treatment restored ejection fraction to levels seen in young controls.

Functionally, the lymphatic inotropic effect of Tf2 translated into enhanced physiological fluid clearance. Using a bioluminescent interstitial clearance assay, Na_V_1.3 activation significantly accelerated third-space fluid drainage, a clinically relevant outcome measure of restoring lymphatic contractility. This supports the idea that pharmacologic enhancement of lymphatic pumping via Na_V_1.3 activation may directly alleviate edema and improve fluid handling in lymphatic disease.

This study has several limitations. The functional experiments were conducted in murine models and utilized pharmacologic agents in acute settings. While human lymphatic tissues demonstrated Na_V_1.3 protein, functional data in human lymphatic vessels is still needed. The finding that Na_V_1.3 is not required for baseline lymphatic contractility may be species-specific to mice. In humans and sheep, baseline lymphatic contractility can be blocked with TTX [44, 45], suggesting that Na_V_1.3 may play a larger role in baseline lymphatic contractility in large animals than in mice. Although Tf2 is highly selective for Na_V_1.3 [59], it is a peptide derived from scorpion venom not yet optimized for human use. Therapeutic development might require modification to optimize pharmacokinetics, immunogenicity, and safety for clinical use, or alternatively the development of a small-molecule isoform-specific activator of Na_V_1.3. In that regard, this study is proof of concept that Na_V_1.3-specific activation has therapeutic potential for lymphatic disease. Chronic studies will also be needed to determine whether repeated or sustained activation of Na_V_1.3 maintains efficacy or induces desensitization or toxicity.

In summary, we exploited Na_V_1.3 as a lymphatic-specific molecular target that regulates lymphatic muscle contractility. Its selective activation with Tf2 reverses dysfunction in aging and radiation injury and promotes interstitial fluid clearance. These findings establish a new pharmacologic strategy to restore lymphatic function in disease.

## ACKNOWLEDGEMENTS

We thank Dr. Peigen Huang and the Cox-7 animal facility for maintaining and providing mice and technical support, Dr. Dai Fukumura and Julia Kahn for maintenance of transgenic mouse lines, Mark Duquette for providing help and technical assistance, and Carolyn Smith for technical assistance with human tissue immunofluorescence staining. We thank Dr. Christopher Lingle (Washington University, St. Louis) for the kind gift of *Scn3a-/-* mice, and Dr. John Wood (University College London) for permission and in whose laboratory this strain originated. We thank Dr. Kiriaki Rajotte for expertise in and editing of MATLAB code for the analysis of lymphatic contractility. We thank the National Institute on Aging at the National Institutes of Health for providing aged mice for this study. Prox1-GFP mice were a kind gift from Dr. Young-Kwon Hong (Beth Israel Deaconess Medical Center, Boston, Massachusetts) and Dr. Taija Makinen (Wihuri Research Institute, Helsinki, Finland). Figures were created in https://BioRender.com.

