## Supplemental Figure 1 for "Selective Activation of Na_V_1.3 Restores Lymphatic Contractility in Age and Injury"

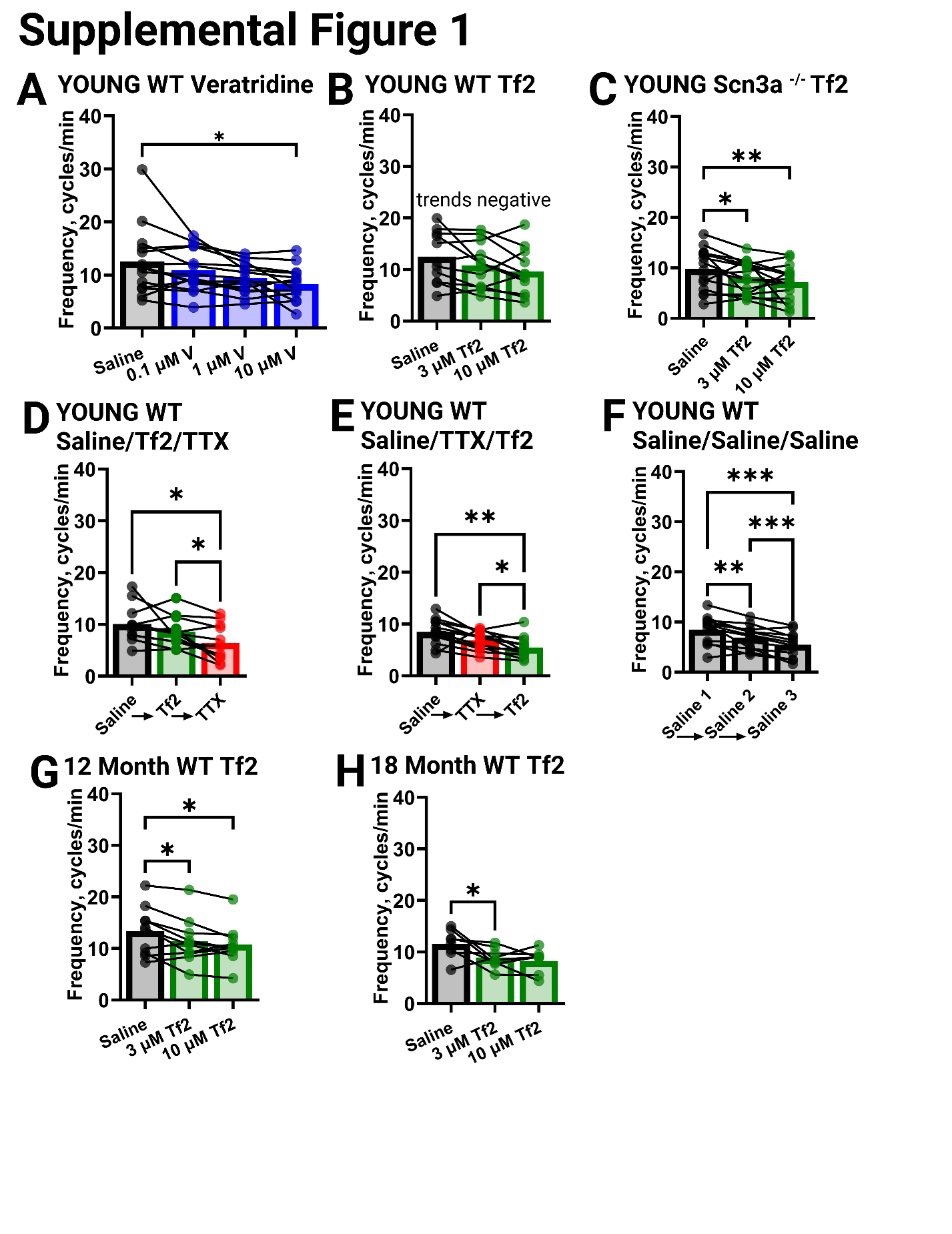

**Supplemental Figure 1:**

The mean frequency of lymphatic contractions measured from data described in Figure 2, Figure 3, and Figure 5 for the following experiments. (A) Young wildtype treated with increasing concentrations of veratridine (V, blue): 0.1 µM, 1 µM, and 10 µM. (B) Young wild type mice treated with increasing concentrations of Tf2 (green): 3 µM, and 10 µM. (C) Young *Scn3a^-/-^* mice treated with increasing concentrations of Tf2 (green): 3 µM, and 10 µM. (D) Young wild type mice treated sequentially with Tf2 (green, 10 µM) then TTX (red, 1 µM ). (E) Young wild type mice treated sequentially with TTX (red, 1 µM ) then Tf2 (green, 10 µM). (F) Young wild type mice sequentially treated with saline (vehicle, black), to show effects over time. (G) 12-month old wild type mice treated with increasing concentrations of Tf2 (green): 3 µM, and 10 µM. (H) 18-month old wild type mice treated with increasing concentrations of Tf2 (green): 3 µM, and 10 µM. Data are from N = 8-16 mice per group. Repeat measures one-way ANOVA with Dunnett’s multiple-comparison test was used for sequential measurements within the same animals with an alpha of 0.05, and adjusted P values with * significance for P<0.03, ** P<0.002, *** P<0.0002.
