## Supplemental Materials and Methods for "Selective Activation of Na_V_1.3 Restores Lymphatic Contractility in Age and Injury"

**METHODS**

**Data Availability Statement**

The original primary data used to generate the figures are available upon reasonable request, pending approval of data transfer agreements between the sending institution (Massachusetts General Hospital) and the receiving institution.

**Animals and Study Design**

Experiments using animals were approved by the MGH Institutional Animal Care and Use Committee (Protocols 2012N000155, 2015N000066, 2024N000028). Studies used 5-12 week old, 12 month old, and 18 month old female and male C57Bl/6 wild-type mice. Several immunofluorescence experiments used C57BL/6 Prox1-GFP mice. 12 month old mice were retired breeders from the Massachusetts General Hospital Cox7 colony, while 18 month old mice were either retired breeders from the Cox7 colony or from the National Institute on Aging at the National Institutes of Health. Prox1-GFP mice were a kind gift from Dr. Young-Kwon Hong (Beth Israel Deaconess Medical Center, Boston, Massachusetts) and Dr. Taija Makinen (Wihuri Research Institute, Helsinki, Finland). Na_V_1.3 knockout mice (*Scn3a-/-)* mice (C57BL/6J background) were kindly provided by Dr. Christofer Lingle (Washington University School of Medicine, St. Louis, MO) with the kind permission of Dr. John Wood (University College London, London, United Kingdom) and maintained as homozygous global knockout mice in our colony [1]. Mice were bred and maintained in the Steele Laboratories for Tumor Biology Cox-7 gnotobiotic animal colony or at the Center for Comparative Medicine at the Charlestown Navy Yard of MGH. Some young C57Bl/6 mice were purchased from Jackson Laboratories.

| **Experimental models: Organisms/strains** | | |
| --- | --- | --- |
| **Strain** | **Source** | **Details** |
| C57BL/6 (5-12 week old) | Massachusetts General Hospital, Cox-7 animal facility | N/A |
| C57Bl/6 (12-month-old) | Massachusetts General Hospital, Cox-7 animal facility, Retired Breeders | N/A |
| C57Bl/6 (18-month-old) | National Institute on Aging | N/A |
| C57Bl/6 (5-12 week old) | The Jackson Laboratories | N/A |
| Prox1-GFP | [2] | RRID: MMRRC_031006-UCD |
| C57Bl/6 *Scn3a-/-* | Massachusetts General Hospital, Cox-7 animal facility | Kind gift from the Christofer Lingle Laboratory, Washington University St. Louis.  Original source John Wood Laboratory, University College London |

***Scn3a-/-* Genotyping Protocol**

Genomic DNA was extracted from mouse tail or ear tissue by proteinase K digestion, precipitation, and resuspension in nuclease-free water. PCR was performed using a three-primer system: a wild-type forward primer, a knockout forward primer, and a common reverse primer. PCR products were amplified with an annealing temperature of 60 °C and resolved on agarose gel electrophoresis. The wild-type allele yields a 212 bp amplicon, while the knockout allele yields a 570 bp amplicon.

| **Genotyping Primers** | |
| --- | --- |
| *Scn3a* wild-type forward primer | 5′- GCTTTTTGTTCAAGTCTATCATATTCAAAG |
| *Scn3a* knockout forward primer | 5′- AAGGATGGCATCACCCACAAG |
| *Scn3a* common reverse primer | 5′-GAGAGAAAGACACTTAAATGCAGACATC |

**Intravital fluorescence lymphangiography**

Mice were anesthetized with an initial subcutaneous injection of ketamine (100-125 mg/kg) and xylazine (10-12.5 mg/kg) and placed on a warming pad. In 30-minute intervals, ketamine (30 mg/kg) was re-dosed if any signs of emergence from anesthesia, such as regaining toe-pinch reflex, were present to maintain a stable plane of anesthesia throughout the study. A 3 μL volume of 2% FITC-Dextran (2 MDa; Millipore Sigma) diluted in 0.9% saline was administered intradermally into the dorsal hind footpad in the second web-space. To expose the afferent lymphatic vessel draining into the popliteal lymph node (PLV), the overlying skin and connective tissue were carefully dissected, following established protocols [3]. Mice were placed sitting upright on a microscope stage taped (silver duct tape, 3M) to a petri dish (100mm x 20mm, Corning 430161) with the experimental hindlimb extended and taped (transpore surgical tape, 3M) lymphatic-side-down against the dish plastic in a custom-built warming water bath and with a custom temperature controller. The temperature of the hindlimb on which the experiment was conducted was measured by an exogenous infrared thermometer gun and maintained between 35-37 °C by adjusting the water bath temperature. The hindlimb was bathed in a 150 µL pool of 0.9% normal saline (Baxter, filter-sterilized daily aliquot) applied to the Petri dish. After undergoing FITC injection, surgery, and placement on the warmed microscope stage on a Petri dish, mice were left to equilibrate and warm for at least 30 minutes on the stage, with periodic evaluation of the lymphatic network contractility. Once stable contractions were robustly present, the solution around the hindlimb was gently removed with a delicate task wiper (Kimwipe) without touching the hindlimb. Fresh 150 µL 0.9% normal saline was applied to the hindlimb, and after 5 minutes of warming and ensuring that the hindlimb was at the correct temperature, recordings were initiated. For pharmacological experiments, all animals served as their own vehicle control. After the baseline recording was made, the 0.9% normal saline was delicately removed with a wipe, and 150 µL containing the test drug at the concentration specified in the data figure and figure legends was applied to the hindlimb. The drug was left to equilibrate for 5 minutes, after which time data was recorded. This same procedure of solution changes and 5-minute incubation was repeated until all paired drug conditions were tested (in this study, between 2-4 conditions).

If a ketamine re-dose was given during equilibration on the microscope, at least 10 minutes passed until recording lymphatic contractility to ensure a stable anesthetic plane during the recording. A stable plane of anesthesia was ensured with no anticipated need for re-dosing ketamine prior to initiating recording a pharmacological experimental series to avoid giving additional anesthetic doses between different drug conditions or altering the experimental time course within a grouped experiment. Time-lapse imaging was conducted on an inverted fluorescence microscope (Olympus IX70), capturing 60-120 seconds of data sampled every 133 ms. Vessel wall motion was analyzed using custom-written MATLAB scripts.

In young mice, ~10% of mice never develop robust lymphatic contractions after undergoing hindlimb surgery to expose the PLV. Since the aim of this study was to measure changes in lymphatic contractility, these mice were excluded from the study and were humanely euthanized. In aged mice, ~20% of mice either never display robust contractility or have FITC uptake in the lymphatic network inadequate to measure an analyzable signal. Though this failure rate in aged mice is decreased by increasing the volume of FITC injected into the hindlimb, we elected to maintain the same volumes of FITC injection between young and old groups as it is known that the volume of fluid injected into the lymphatic network impacts the contractile properties of the lymphatic vessels [4].

**Analysis of Lymphatic Contractility**

Custom-written code (MATLAB R2023a, MathWorks, Inc.) was used for image analysis to calculate the lymphangion ejection fraction, frequency, mean vessel diameter, systolic (‘valley’) diameter, diastolic (‘peak’) diameter. After importing the tiff data files and the frame rate, the Image Processing Toolbox imbinarize function is used with a fixed global threshold to binarize the grayscale image into one that is black and white. The diameter of each vessel in each tiff is then calculated. This value is plotted over time to create a classic peak-and-valley plot (peaks corresponding to diastole, and valleys corresponding to systole of the lymphangion pumping). The mean vessel diameter for each recording is calculated as the arithmetic mean of the diameter over the entire recording. The systolic vessel diameter is the average of all diameter ‘valley’ values, while the diastolic vessel diameter is the average of all diameter ‘peak’ values. The frequency is calculated as the number of cycles (peak followed by a valley) in the recording per minute. The ejection fraction of a single lymphatic contraction is calculated by assuming that the two-dimensional vessel image corresponds to a cylindrical vessel, and projected area of the vessel is transformed into volume by $Volume=\pi{(\frac{Diameter}{2})}^{2}x Length$. The ejection fraction of a single contraction is ${(V_{D}-V_{S})}/{V_{D}}$, where V_D_ is the diastolic (‘peak’) volume, and V_S_ is the systolic (‘valley’) volume. The single ejection fraction for a recording is the average ejection fraction over all contraction cycles in that recording.

**Bioluminescence Interstitial Fluid Clearance Assay**

Bioluminescence imaging was conducted using the IVIS Spectrum System (PerkinElmer) and controlled with Living Image software (version 4.3.1). The “Luminescent” and Photograph” settings were selected under Imaging Mode within the IVIS Acquisition Control Panel. The stage temperature was set to 35°C and confirmed to be within 35-37°C using an exogenous infrared thermometer. The exposure time was set to 90 seconds and saved to “Seq-1.” A trial recording without mice was performed at the beginning of each session to ensure proper lens focus.

Mice were anesthetized with an initial subcutaneous injection of ketamine (100-125 mg/kg) and xylazine (10-12.5 mg/kg), then placed on a warming pad. The lower ventral surface and both hindlimbs were disinfected with 70% ethanol, and 500 µL of 30 mg/mL D-Luciferin (PerkinElmer) was administered intraperitoneally (IP) in the left lower quadrant. To expose the afferent lymphatic vessel draining into the popliteal lymph node (PLV), the overlying skin and connective tissue were carefully dissected, following established protocols [3], ensuring preservation of the ventral hindlimb skin for imaging.

Mice were positioned supine on a light-controlled black aluminum sheet (Rosco Blackwrap), and hindlimbs were extended and secured with transpore surgical tape (3M), with the lymphatic side facing downward against the light-controlled sheet. Hindlimb temperatures were maintained between 35-37°C and confirmed using an exogenous infrared thermometer gun. Each hindlimb was bathed in a 150 µL pool of 0.9% normal saline (Baxter, filter-sterilized daily aliquot). After 25 minutes, 2 μL of luciferase (5 mg/mL; Perkin Elmer) was injected intradermally into the dorsal hind footpad (second web-space) of both hindlimbs using two injector tools (one per footpad) connected to 30G needles, 3” in length PE-10 tubing, and 1 mL syringes. 500 μL of filter-sterilized 0.9% normal saline was injected IP into the right lower quadrant for hydration, and the mouse was carefully placed into the whole-body imaging chamber.

Images were acquired every 150 seconds. For pharmacological experiments, all animals served as their own vehicle control. Following a 15-minute baseline recording, the bathing solution around each hindlimb was removed using a delicate task wiper (Kimwipe), avoiding direct contact with the exposed hindlimb. Fresh 150 μL of 0.9% normal saline was applied to one hindlimb (negative control), while 150 µL containing the test drug was applied topically to the contralateral hindlimb dropwise using a P200 pipette. To control for potential laterization bias, the assignment of experimental treatment to the left or right hindlimb was alternated across the cohort. Imaging continued at 150-second intervals for the duration of the experiment.

Ketamine (30 mg/kg) was re-administered subcutaneously every 60 minutes, or as needed, based on signs of anesthesia emergence (e.g., toe-pinch reflex), to maintain a stable plane of anesthesia throughout the study. The imaging chamber was opened every 30 minutes for animal wellness checks, including confirmation of body temperature (35-37°C), and to ensure proper ventilation. To prevent dehydration, 500 μL of filter-sterilized 0.9% normal saline was injected IP into the right lower quadrant every 60 minutes.

Bioluminescence interstitial fluid clearance was analyzed using custom-written two-compartment model MATLAB scripts.

**Analysis of Bioluminescence Clearance Kinetics**

Regions of interest were defined bilaterally at the center of the footpads immediately following acquisition of the first image. Each ROI was standardized to a fixed size of 17x17 pixels and applied in an automated manner to all subsequent measurements for each mouse throughout the duration of the experiment. Quantitative outputs from each ROI were manually recorded into a Microsoft Excel spreadsheet, with values reported to four significant figures. The completed spreadsheet was then imported into MATLAB for quantitative analysis using a two-compartment model.

To quantify clearance kinetics from the footpad following luciferase injection, we applied a two-compartment kinetic model with a single absorption site (Compartment 1: interstitial footpad tissue) and a clearance compartment (Compartment 2: lymphatic drainage or systemic clearance). The model assumes first-order kinetics for both absorption into the first compartment and elimination from the second compartment. A(t) represents the mass of signal remaining at the injection site (Compartment 1) and C(t) represents the concentration of signal in the systemic clearance compartment (Compartment 2). The system is governed by: $\frac{dA}{dt}= -k_{a}A(t)$ and $\frac{dC}{dt}= k_{a}A\left( t \right)- k_{e}C\left( t \right)$ with initial conditions $A\left( 0 \right)= A_{0}$, $C\left( 0 \right)= 0$. The solution for the concentration in the second compartment (C) following a bolus input at time zero is: $C\left( t \right)= A_{0}\frac{k_{a}}{k_{a}-k_{e}}(e^{-k_{e}t}-e^{-k_{a}t})$. Fitting was performed in MATLAB (MATLAB R2023b) using the fminsearch function to minimize the sum of squared residuals between the model and experimental data. The fitted curve was plotted against experimental data to assess the goodness of fit. K_e_ values were reported in the final results. Data from several recordings showed only the decay curve phase rather than both an uptake and decay and did not fit 2-compartment modeling. In these cases, k_e_ was measured in Excel using the built-in exponential decay trendline function. Goodness of fit was evaluated visually.

**Radiation-Induced Lymphatic Damage Model**

Male and female C57BL/6J mice (7–10 weeks old) were randomly assigned to either a radiation treatment or a sham treatment control group. In the radiation group, the right hindlimb of each mouse was selectively exposed to X-ray radiation to assess the localized effects of acute radiation on peripheral lymphatic vessels. The rest of the body was shielded from radiation exposure using custom-designed lead protectors. Before each irradiation session, mice were anesthetized with subcutaneous injection of ketamine (125 mg/kg) and xylazine (12.5 mg/kg) and placed on a warming pad. A total dose of 18 Gy was administered, delivered as 6 Gy per day over three consecutive days. Irradiation was performed using a Precision X-ray irradiator operating at 225 kVp and 13.30 mA, with a dose rate of 1.98 Gy/min. Sham-treated mice underwent identical handling and anesthesia protocols, but were not exposed to radiation.

**Human lymphatic vessel tissue**

Human subjects were consented under Beth Israel Deaconess Medical Center (BIDMC) IRB# 2017P000190, “Lymphedema Surgery Biorepository.” The samples were transferred to MGH via protocol BIDMC IRB 2018P000191, “An Evaluation of Lymphatic Surgery Samples”, with an MTA in place between the Padera lab at MGH and the Singhal lab at BIDMC. Surgical samples were obtained by Dr. Dhruv Singhal during axillary lymph node dissection (ALND) for the treatment of breast cancer followed by immediate lymphatic reconstruction (ILR). During the ILR, free ends of lymphatic vessels are collected that otherwise would have been discarded. Three samples were collected with demographics in the table below. Samples were placed on ice, then preserved in OCT (tissue tek) and cooled using an ethanol slurry within 10 minutes of removal. They were then stored at -80°C until used later for immunofluorescence.

| **Human Lymphatic Vessel Sample Information** | | | |
| --- | --- | --- | --- |
| Sample Number | Sex | Race | Age Group |
| 209 | Female | Non-Hispanic White | 60-69 |
| 220 | Female | Non-Hispanic White | 40-49 |
| 262 | Female | Non-Hispanic White | 70-79 |

**Collecting lymphatic vessel isolation**

Inguinal-axillary mouse lymphatic collecting vessels were surgically isolated and collected for qPCR and immunofluorescence whole-mount studies as previously described [5]. In brief, mice were anesthetized by subcutaneous injection of ketamine (100 mg/kg) and xylazine (10 mg/kg). Fur on the entire ventral surface was shaved, including the forelimbs and hindlimbs, and fur debris was removed with delicate task wipers and adhesive tape. The surgical area was sterilized with 70% ethanol. A midline incision was made along the ventral abdomen, extending from the pubis to the sternum, with additional lateral cuts made to the middle of all four limbs. The skin and subcutaneous tissue were peeled laterally using blunt dissection, leaving the peritoneum intact. This exposes the inguinal-axillary collecting lymphatic vessels (IALV) that run between the inguinal and axillary lymph nodes. To visualize the lymphatic network, 10 μL of 0.5% Evans Blue dye in PBS was injected directly into the inguinal lymph node, allowing the dye to fill the lymphatic vessels to the axillary lymph node. Under a stereomicroscope, the blue collecting lymphatics were carefully dissected in situ. Surrounding fat and connective tissue were removed while the vessels were still connected to the nodes. The lymphatics are in a unique tissue plane, and with gentle tension can glide out from surrounding tissues and be isolated in situ. Once the entire clean vessel is exposed, it is cut at the inguinal and axillary nodes and removed from the mouse for further processing. Typically 3-6 cm of vessel per side are retrieved. If additional anesthesia is needed during the procedure, doses of 30 mg/kg ketamine are administered.

**Immunofluorescence**

**Mouse**

Murine inguinal axillary lymphatic collecting vessels were isolated as described above. Vessels were fixed for 30 minutes at room temperature in 4% formaldehyde and washed three times in 1X PBS for 15 minutes at room temperature. Vessels were then incubated in blocking/permeabilization buffer containing 10% normal donkey serum and 0.1% Triton X-100 for three hours at room temperature. Vessels were then washed three times in 1X PBS for 5 minutes and cut into 5-10 mm pieces and placed into individual wells of a 96-well plate. For primary antibody staining, each vessel piece was incubated overnight at 4°C with aSMA-Cy3 (1:200 dilution, Millipore Sigma, Cat #: C6198), DAPI (1:1000 dilution, Invitrogen, Cat #: D1306), and Anti-Na_V_1.3 antibody (1:125 - 1:250 dilution, Alomone, Cat #: ASC-023), prepared in 1% normal donkey serum and 0.05% Triton X-100. After primary antibody incubation, vessel pieces underwent three washes in 1X PBS for 15 minutes each at room temperature, followed by overnight incubation at 4°C with Alexa Fluor 647 donkey anti-rabbit secondary antibody (1:200 dilution, Jackson ImmunoResearch, Cat #: 711-605-152) prepared in 1% normal donkey serum. Upon completing this step, three washes were performed in 1X PBS for 15 minutes each at room temperature. Vessel pieces were mounted in straight lines on microscope slides with Vectorshield Plus mounting medium (Vector Laboratories, Cat #: H-1900). Lymphatic vessels were imaged with a Zeiss LSM 880 Superresolution (Airyscan) microscope.

**Human**

Tissue blocks were sliced with a cryostat and placed directly onto slides. Slides were air-dried for 10 min before starting the staining protocol. Slides were post-fixed in acetone at -20°C for 5 min, washed in PBS, blocked with 5% Normal Donkey Serum in PBS, and incubated with anti-Na_V_1.3 antibody (Alomone ASC-023) overnight at 1:250 dilution. The next day, they were washed and incubated with secondary antibody (Jackson ImmunoResearch Alexa 488 anti-rabbit secondary) (1:200), anti-αSMA-Cy3 (Sigma 6198) (1:200), and DAPI for 2 hours at room temperature. Slides were then washed with PBS and cover slips were mounted using Vectashield Vibrance cover slipping medium (Vector Labs).

| **Antibodies** | | |
| --- | --- | --- |
| **Reagent** | **Source** | **Identifier** |
| Anti-NaV1.3 antibody (external epitope), Rabbit host | Alomone | ASC-023 |
| Anti-actin, α-Smooth muscle-Cy3 antibody, Mouse monoclonal | Millipore Sigma | Cat#: C6198; RRID: AB_476856 |
| Anti-Rabbit IgG secondary antibody, Alexa Fluor 647 | Jackson ImmunoResearch | 711-605-152 |

**RNA extraction and real-time quantitative PCR assay**

Following inguinal axillary lymphatic vessel collection as described above, vessels were immediately transferred into 1.5 mL cryogenic vials (ThermoFisher Scientific) and snap-frozen in liquid nitrogen to maintain RNA integrity. Total RNA was extracted using the PicoPure RNA Isolation Kit (ThermoFisher Scientific, Cat. #: KIT0202, KIT0204) following the manufacturer’s protocol. Complementary DNA (cDNA) was synthesized using the iScript cDNA Synthesis Kit (Bio-Rad) according to standard procedures. Each RNA extraction was performed using four inguinal axillary lymphatic vessels pooled from two mice. Resulting cDNA samples from multiple extractions were combined into a single master tube. Due to expected low absolute RNA yields in each sample, cDNA was diluted 1:3 for PCR reactions. Quantitative real-time PCR was then carried out for selected target genes using iTaq Universal SYBR Green Supermix (Bio-Rad, Cat. #: 1725124) on a Stratagene MxPro 30005 Real-Time PCR Detection System (Agilent), following the manufacturer’s guidelines.

**Primer Sequences**

| **Gene** | **Forward Primer** | **Reverse Primer** |
| --- | --- | --- |
| *Gapdh (mouse)* | 5’ AGG TCG GTG TGA ACG GAT TTG 3’ | 5’ TGT AGA CCA TGT AGT TGA GGT CA 3’ |
| *Scn3a (mouse)* | 5’ TGT GGG ACT GCA TGG AGG TC 3’ | 5’ CAT CGT CCG TAG CAG CAA GG 3’ |

**Single Cell Sequencing Data Analysis**

Murine lymphatic vessel single-cell RNA-seq datasets (GSE248658 and GSE277843) were downloaded from the NCBI GEO database. Single-cell RNA-seq data were analyzed using Seurat (version: 5.1.0), with “SCTransform” applied for normalization and data scaling and the top 30 principal components used for Uniform Manifold Approximation and Projection (UMAP) analysis. Cell type annotation was performed using both SingleR (version: 2.6.0) and the top highly expressed genes in each cluster identified. After identifying lymphatic muscle cells, these cells were integrated using the 'reciprocal PCA' algorithm in Seurat. Data visualization was performed using SeuratExtend (version: 1.1.4).

**Statistics**

Data are expressed as mean ± SEM. Repeat measures one-way ANOVA with Dunnett’s multiple-comparison test was used for sequential measurements within the same animals with an alpha of 0.05, and adjusted P values with * significance for P<0.03, ** P<0.002, *** P<0.0002. Unpaired t-tests were used for comparisons between independent groups with the same significance cutoffs as above. Comparisons within the same animal were performed using a two-tailed paired t-test, with * indicating P<0.05. Analyses were conducted in GraphPad Prism.
